# Tissue-Specific Dynamics of *TCF4* Triplet Repeat Instability Revealed by Optical Genome Mapping

**DOI:** 10.1101/2024.03.27.587034

**Authors:** Christina Zarouchlioti, Stephanie Efthymiou, Stefano Fracchini, Natalia Dominik, Nihar Bhattacharyya, Siyin Liu, Marcos Abreu Costa, Anita Szabo, Amanda N Sadan, Albert S Jun, Enrico Bugiardini, Henry Houlden, Andrea Cortese, Pavlina Skalicka, Lubica Dudakova, Kirithika Muthusamy, Micheal E Cheetham, Alison J Hardcastle, Petra Liskova, Stephen J Tuft, Alice E Davidson

## Abstract

Here, we demonstrate the utility of optical genome mapping (OGM) to interrogate the Fuchs endothelial corneal dystrophy (FECD)-associated intronic TCF4 triplet repeat (termed CTG18.1) and gain novel insights into the tissue-specific nature of the disease. Genomic DNA (gDNA) samples derived from peripheral blood leukocytes and primary corneal endothelial cells (CECs) were analysed by OGM. Concurrently, all samples were genotyped by standard PCR-based methods to classify their expansion status. Individuals with one or more CTG18.1-expanded alleles (≥50 CTG repeats) detected in their leukocyte-derived gDNA were classified as expansion-positive. A customised bioinformatics pipeline was developed to perform CTG18.1-targeted OGM analysis. All linearised gDNA molecules containing labels flanking CTG18.1 were extracted, corrected for the repeats on the reference human genome and sized. Analysis of paired bio-samples revealed that expanded CTG18.1 alleles behave dynamically, regardless of cell-type origin, but displayed significantly higher levels of instability within the diseased corneal endothelium. Clusters of CTG18.1 molecules of approximately 1,800-11,900 repeats, beyond the ranges observed in individual-matched leukocyte samples, were detected in all CEC gDNA samples from expansion-positive cases. In conclusion, OGM is a powerful method to analyse the somatically unstable CTG18.1 locus. More generally, this work exemplifies the broader utility of OGM in exploring somatically unstable short tandem repeat loci. Furthermore, this study has highlighted the extreme levels of tissue-specific CTG18.1 somatic instability occurring within the diseased corneal endothelium, which we hypothesise plays a pivotal role in driving downstream pathogenic mechanisms of CTG18.1-mediated FECD.

## Introduction

Fuchs endothelial corneal dystrophy (FECD) is a common age-related cause of heritable visual loss primarily affecting the corneal endothelium (Matthaei et al. 2019). The post-mitotic monolayer of corneal endothelial cells (CECs) maintains a leaky barrier that regulates corneal hydration (Joyce 2012). In FECD, accelerated CEC loss occurs, compromising barrier function and progressively resulting in corneal swelling, clouding and reduced visual acuity (Price et al. 2021). To date, expansion of an intronic triplet repeat within the *TCF4* gene (termed CTG18.1; OMIM #613267) has been identified as the most common risk factor for FECD in all ethnic groups studied (Fautsch et al. 2021). Remarkably, such expansions are detected in up to 81% of European FECD patient cohorts, making it by far the most common short tandem repeat (STR) expansion disease in humans (Zarouchlioti et al. 2018; Fautsch et al. 2021). Thus, as well as being a common cause of age-related vision loss, FECD represents an important paradigm for much rarer, currently incurable and devastating STR diseases, such as Huntington’s disease (OMIM #143100) and myotonic dystrophy (OMIM #160900; 602668).

STRs are somatically unstable elements that can contract and expand in an age-, repeat length- and tissue-specific manner (Mouro Pinto et al. 2020; Wheeler and Dion 2021; Monckton 2021). We have previously utilised a long-read amplification-free sequencing method to determine that expanded copies of CTG18.1 (defined as ≥50 repeats) are somatically unstable in peripheral blood leukocytes, with inherited repeat lengths positively correlating with increased CTG18.1 instability (Hafford-Tear et al. 2019). Southern blot data of FECD patient-derived corneal endothelial cell cultures have also suggested that CTG18.1 repeat length is greater in affected cells compared to leukocytes, in keeping with the dogma that STRs typically display high levels of instability within affected cell types (Wieben et al. 2021; Depienne and Mandel 2021). For example, in myotonic dystrophy type 1, the disease-associated repeat has been reported to expand up to 4,000 repeats in skeletal muscle, representing up to a 25-fold increase in length compared to blood (Nakamori et al. 2013). Multi-omic data studies of Huntington’s disease have recently brought disease-associated STR instability mechanisms to the forefront of translational genomic medicine, revealing that repeat-length mosaicism within affected cell populations is a fundamental driver of disease (Handsaker et al. 2023).

Disease-associated STR instability is hypothesised to be a unifying mechanism and translationally relevant pathway for many STR-associated diseases (Benn et al. 2021). However, it can be extremely challenging to measure repeat length mosaicism in affected cell types, given the rarity of relevant material and the technical difficulties associated with accurately sizing or sequencing large repetitive and unstable genomic elements. Conventionally, STR length is assessed by gel or capillary electrophoresis of PCR amplicons and/or Southern blot (Ciosi et al. 2021). More recently, short-read next-generation sequencing (NGS) approaches have been developed to quantify levels of repeat length instability (Ciosi et al. 2019). Amplification-free long-read sequencing methods have also yielded novel insights but require high levels of input DNA (Erdmann et al. 2023; Höijer et al. 2018). Thus, collectively, these approaches are either restricted by repeat size detection thresholds or the large inputs of DNA required, which are scarcely available from affected patient-derived tissues.

Optical Genome Mapping (OGM) offers a powerful method to interrogate native ultra-long molecules of DNA (>150 kb) to reveal large and/or complex structural variants (>500 nucleotides) across the genome (Yuan et al. 2020). To date, it has been suggested as a viable way to detect STR expansions and contractions associated with a subset of neurological and neuromuscular conditions (Facchini et al. 2023; Morato Torres et al. 2022; Maroilley et al. 2023; Efthymiou et al. 2023). Despite *TCF4* being ubiquitously expressed, CTG18.1 expansions have only been robustly associated with FECD, a corneal endothelial cell-specific disease. This knowledge, in combination with increased levels of somatic instability reported in other disease-associated STR loci within affected cell populations, prompted us to explore CTG18.1 length and instability in patient-derived CECs.

Here, for the first time, we apply OGM to enable single molecule resolution of CTG18.1 in a series of genomic DNA samples from expansion-positive individuals affected by FECD. Utilising gDNA samples isolated from unaffected peripheral blood leukocytes and affected corneal endothelial cells, we explore the utility of this method to size the repeat and, importantly, to gain novel insights into the tissue-specific nature of this common and sight-threatening disease.

## Results

### Application of OGM to interrogate CTG18.1 repeat expansions

CTG18.1 is typically sized from peripheral blood leukocyte gDNA samples using a dual PCR-based approach that detects and sizes PCR amplicons by capillary electrophoresis (Wieben et al. 2012; Mootha et al. 2014). This protocol consists of an STR PCR-based assay to size CTG18.1 alleles of ≤120 CTG repeats (**Figure 1A** i,iii) and a triplet-primed PCR assay (TP-PCR) to detect the presence or absence of expanded alleles, including those beyond the detection limit of the STR-PCR assay (>120 repeats) (**Figure 1A** ii,iv,vi). Despite proving an efficient and precise way to genotype leukocyte-derived CTG18.1 alleles, the upper detection limit of the STR assay precluded sizing of 3.3% (14/427) expanded CTG18.1 alleles reported in our FECD patient DNA bio-resource (n=450) (Zarouchlioti et al. 2018).

**Figure 1.**
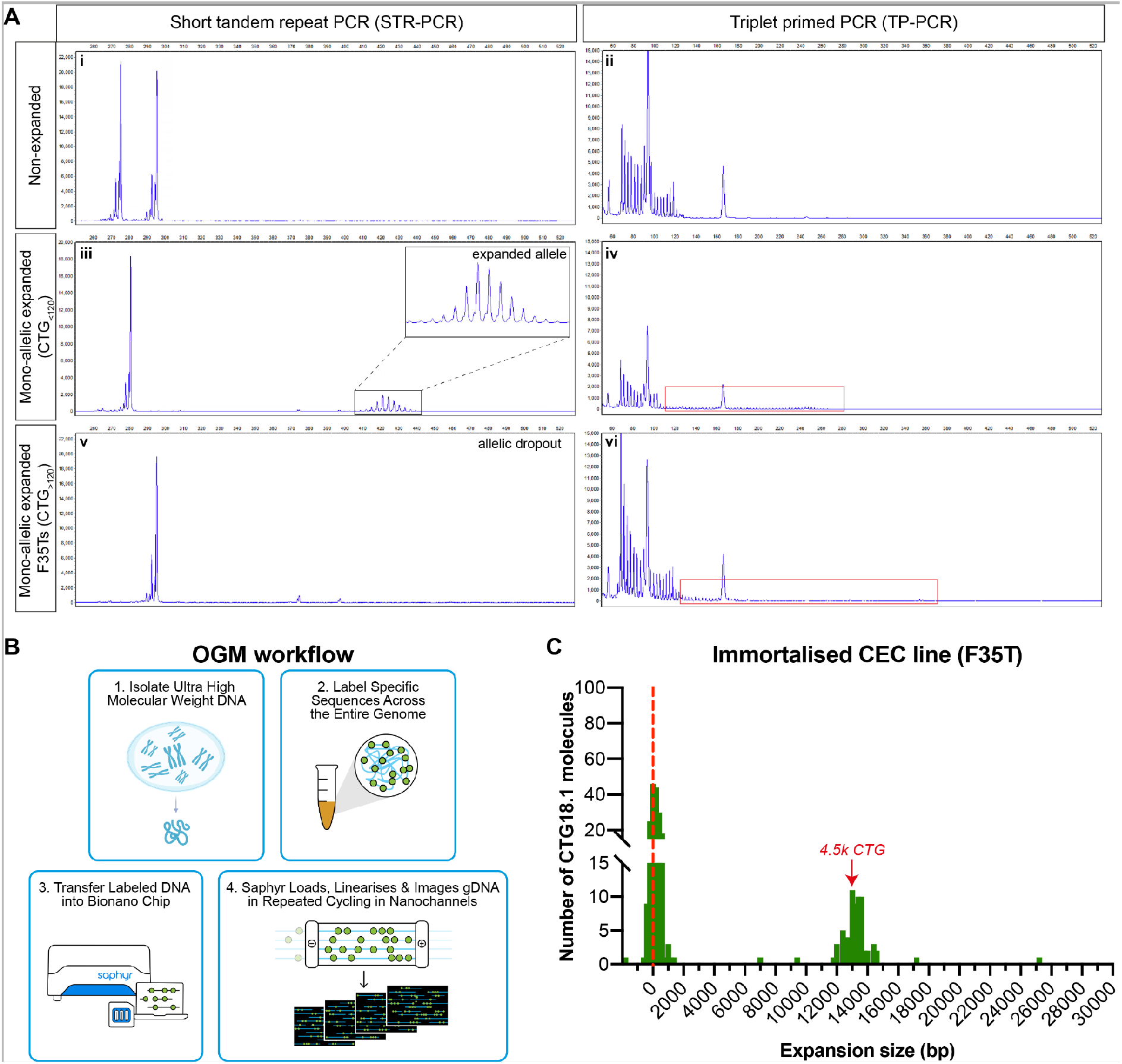
Optical genome mapping (OGM) effectively detects large expanded CTG18.1 alleles. **(A)** Detected traces after capillary electrophoresis of STR-PCR (i, iii, v) and TP-PCR (ii, iv, vi) products amplified from non-expanded whole-blood derived gDNA samples (i-ii), mono-allelic expanded whole-blood derived (iii-iv) and F35T cell-derived (v-vi) gDNA samples. Red boxes highlight the presence of expanded alleles as indicated by TP-PCR traces. **(B)** Schematic summary of OGM methodology; (1) extraction of ultra-high molecular weight (UHMW) gDNA that is (2) subsequently labelled via covalent modification at genome-wide CTTAAG hexamer motifs before (3) linearising and imaging the decorated molecules on nanochannels (image adapted from: https://bionanogenomics.com). (**C**) Histogram of OGM CTG18.1 molecule sizes (bp) observed in immortalised CEC line F35T. The red dotted line indicates alleles around the lowest detection threshold of the method, likely representing the non-expanded allele.

Given that OGM can detect very large structural variants, we aimed to investigate the utility of this method to interrogate CTG18.1 and explore the capabilities of this assay to reveal ultra-long CTG18.1 expanded alleles. As a positive control, we selected the F35T immortalised corneal endothelial cell line as it has previously been reported by Southern blot to have a monoallelic expansion of ∼4,500 CTG repeats (Saha et al. 2022). When analysed with the STR assay, only one CTG18.1 allele harbouring 21 CTG repeats was detected (i.e., allelic dropout is observed). In contrast, TP-PCR confirmed the presence of a longer expanded allele beyond the detection threshold of the STR assay **(Figure 1A** v-vi).

Ultra-high molecular weight (UHMW) gDNA was extracted from F35T cells and analysed by OGM (**Figure 1B**). After correcting for the flanking region between the labels of interest and the 24 CTG repeats in hg38, the size of the CTG18.1 interval was determined for each individual DNA molecule (**Figure 1C**). An accumulation of molecules was detected around 0 bp, likely representing the shorter non-expanded allele, as this is below the lower detection limit of this assay (±500 bp). Importantly, a cluster of 13,500 bp molecules was also detected, corresponding to approximately 4,500 CTG repeats, replicating previously reported Southern blot data (Saha et al. 2022). A small number of additional DNA molecules was detected above (up to 25.2 kb long) and below (single molecule at -1.8 kb) the expected size range, based on the dual PCR assay and Southern blot data. These single non-clustering molecules were deemed likely to be artefacts arising from the misalignment of long repetitive regions with relatively sparse fluorescent labels (Mantere et al. 2021). Nonetheless, the detection of molecules clustering at approximately 13,600 bp (representing 4,500 repeats) indicated the presence of a large CTG18.1 allele that evaded sizing by the dual PCR-based genotyping assay and was concordant with Southern blot data, thus supporting the utility of OGM to detect the presence of large expanded CTG18.1 alleles (**Table 1**).

**Table 1.** Summary of subjects and gDNA samples analysed by optical genome mapping. Karyotypic sex, diagnosis, age of first corneal transplant surgery (as an indicator for disease severity), and sampling age summarised for all subjects and samples of the study. STR: short-tandem repeat genotyping, BL: peripheral blood leukocyte sample, CEC: corneal endothelial cell sample, N/A: not applicable information. TP-PCR + and - indicates the presence or absence of an expanded allele. Subject sampling age shown for both blood collection and excised corneal endothelial tissue. *Genotyping data is derived from a patient-matched leukocyte gDNA sample.

| Subject | Sex | FECD<br>Diagnosis | Age of 1<sup>st</sup><br>keratoplasty | STR<br>genotype | TP-PCR | Subject<br>sampling age | Sample<br>type | OGM Mean<br>allele<br>length<br>(bp) | OGM Max<br>allele<br>length (bp) | Number of<br>CTG18.1<br>molecules |
| --- | --- | --- | --- | --- | --- | --- | --- | --- | --- | --- |
| **Immortalised corneal endothelial cell (CEC) line** |  |  |  |  |  |  |  |  |  |  |
| F35T | Female | ✓ |  | 21/ | + | N/A |  | 10,079 | 25,159 | 250 |
| **Patient-matched peripheral blood leukocytes (BL) and primary CEC-derived gDNA sample series** |  |  |  |  |  |  |  |  |  |  |
| 1 | Male | ✓ | 61 | 16/63<br>16/ | +<br>+ | 62<br>62 | BL-1<br>CEC-1 | 196<br>2,771 | 1,849<br>1,7798 | 438<br>254 |
| 2 | Female | ✓ | 59 | 17/63<br>17/ | +<br>+ | 59<br>59 | BL-2<br>CEC-2 | 162<br>2,752 | 1,162<br>19,758 | 328<br>251 |
| 3 | Female | ✓ | 75 | 31/75<br>31/ | +<br>+ | 76<br>77 | BL-3<br>CEC-3 | 200<br>2,946 | 1,345<br>27,124 | 372<br>245 |
| 4 | Male | ✓ | 64 | 11/77<br>11/ | +<br>+ | 64<br>64 | BL-4<br>CEC-4 | 216<br>2,192 | 1,033<br>13,247 | 365<br>358 |
| 5 | Male | ✓ | 83 | 27/81<br>27/ | +<br>+ | 83<br>83 | BL-5<br>CEC-5 | 371<br>2,827 | 2,129<br>32,265 | 472<br>246 |
| 6 | Female | ✓ | 69 | 26/85<br>26/ | +<br>+ | 69<br>69 | BL-6<br>CEC-6 | 282<br>3,625 | 1,484<br>35,561 | 396<br>266 |
| 7 | Female | ✓ | 59 | 11/90<br>11/ | +<br>+ | 53<br>59 | BL-7<br>CEC-7 | 303<br>1,977 | 1,920<br>15,824 | 400<br>279 |
| 8 | Female | ✓ | 65 | 11/90<br>11/ | +<br>+ | 65<br>66 | BL-8<br>CEC-8 | 481<br>2,101 | 5,437<br>19,513 | 373<br>247 |
| 9 | Male | ✓ | 58 | 25/96<br>25 | +<br>+ | 58<br>58 | BL-9<br>CEC-9 | 408<br>2,698 | 4,670<br>19,924 | 308<br>286 |
| **CTG18.1 expansion-positive peripheral blood leukocytes (BL)-derived gDNA sample series** |  |  |  |  |  |  |  |  |  |  |
| 10 | Male | ✓ | 70 | 64/82 | + | 72 | BL-10 | 383 | 1,890 | 432 |
| 11 | Male | ✓ | 77 | 11/85 | + | 69 | BL-11 | 291 | 3,113 | 358 |
| 12 | Male | ✓ | 54 | 11/94 | + | 54 | BL-12 | 334 | 1,302 | 376 |
| 13 | Female | ✓ | 65 | 68/95 | + | 73 | BL-13 | 590 | 7,055 | 360 |
| 14 | Male | ✓ | 74 | 11/101 | + | 74 | BL-14 | 1039 | 8,412 | 297 |
| 15 | Male | ✓ | 67 | 17/107 | + | 67 | BL-15 | 996 | 9,179 | 336 |
| **CTG18.1 expansion-negative peripheral blood leukocyte (BL-Ctr)-derived gDNA sample series** |  |  |  |  |  |  |  |  |  |  |
| 16 | Female | ✗ | N/A | 11/11 | - | 57 | BL-Ctr1 | 172 | 1,385 | 509 |
| 17 | Female | ✗ | N/A | 11/16 | - | 29 | BL-Ctr2 | 117 | 1,140 | 396 |
| 18 | Female | ✗ | N/A | 11/16 | - | 50 | BL-Ctr3 | 110 | 1,265 | 391 |
| 19 | Male | ✓ | 92 | 14/21 | - | 92 | BL-Ctr4 | 109 | 1,258 | 415 |
| 20 | Male | ✗ | N/A | 11/22 | - | 22 | BL-Ctr5 | 134 | 1,326 | 309 |
| 21 | Male | ✗ | N/A | 17/22 | - | 40 | BL-Ctr6 | 135 | 994 | 363 |
| 22 | Male | ✗ | N/A | 17/23 | - | 67 | BL-Ctr7 | 167 | 1,174 | 396 |
| 23 | Male | ✗ | N/A | 23/24 | - | 48 | BL-Ctr8 | 184 | 1,386 | 414 |
| 24 | Female | ✗ | N/A | 17/32 | - | 69 | BL-Ctr9 | 142 | 1,450 | 407 |
| **CTG18.1 expansion-negative corneal endothelial cell (CEC)-derived gDNA sample series** |  |  |  |  |  |  |  |  |  |  |
| 25 | Female | ✓ | 60 | 11/14 | - | 60 | CEC-Ctr1 | 111 | 1,340 | 470 |
| 26 | Female | ✗ | N/A | 11/17 | - | 54 | CEC-Ctr2 | 51 | 1,281 | 369 |
| 27 | Male | ✗ | N/A | 15/17 | - | 70 | CEC-Ctr3 | 164 | 1,751 | 399 |
| 28 | Female | ✗ | N/A | 25/27 | - | 50 | CEC-Ctr4 | 110 | 904 | 198 |
| 29 | Male | ✗ | N/A | 11/32 | - | 74 | CEC-Ctr5 | 97 | 979 | 351 |
| 30 | Male | ✗ | N/A | 11/39 | - | 58 | CEC-Ctr6 | 182 | 1,087 | 415 |
| **CTG18.1 bi-allelic corneal endothelial cell (CEC)-derived gDNA sample** |  |  |  |  |  |  |  |  |  |  |
| 31 | Female | ✓ |  | 57/93* |  | 77 | CEC-10 | 16,803 | 30,232 | 87 |

### CTG18.1 displays tissue-specific somatic instability

In the knowledge that OGM can effectively detect and size a large expanded CTG18.1 allele in the immortalised F35T cell line, we next wanted to explore CTG18.1 length and instability levels in the primarily affected cell population. Given that patients with FECD typically only undergo corneal transplantation when there is advanced disease, when the CEC density has substantially declined, total cell number and gDNA yields from these specimens are extremely low. To overcome this limitation and acquire enough cells for the OGM protocol (approximately 1 million cells per sample), we used primary CEC cultures generated from endothelial keratoplasty specimens removed following planned corneal surgery to analyse CEC-specific gDNA. We have previously demonstrated that these primary CEC cultures robustly maintain the biomarkers of CTG18.1-mediated disease and generate a pure corneal endothelial cell model that enables the investigation of FECD in a disease-relevant cell context (Zarouchlioti et al. 2018; Bhattacharyya et al. 2024).

We acquired paired biosamples (corneal endothelial specimens and peripheral blood leukocytes) from 9 FECD patients to directly compare instability levels between the affected and unaffected cell types. Samples were selected to reflect the overall range of expanded alleles observed in our patient cohort (Zarouchlioti et al. 2018). UHMW genomic DNA from the paired leukocytes and CEC samples was isolated using the recommended Bionano protocols. Initially, all samples were analysed using the dual PCR-based genotyping method (**Table 1**). For all leukocyte-derived samples, two alleles were detected, one non-expanded and one expanded (<120 CTG repeats) allele. However, similar to the F35Ts, only one non-expanded allele was detected by STR-PCR in all paired CEC samples. Thus, allelic dropout of larger alleles beyond the sizing threshold of the STR assay consistently occurred, while TP-PCR indicated the presence of an expanded allele in all expansion-positive samples (**Table 1**), as previously observed with the F35T cell line (**Figure 1A**).

Next, the paired gDNA samples were analysed by OGM (**Figure 1B; 2**). The mean detectable repeat size in the leukocyte-derived gDNA sample series was approximately 97 CTG repeats (range 54-160) (**Figure 2**). These figures reflect the number of molecules detected from the expanded and non-expanded alleles (unphased molecules) within each of the nine samples. As most of the molecules from the respective samples are below the lower detection threshold of the assay (500 bp) they could not be sized with confidence. However, when comparing the mean molecule size of each of the nine leukocyte samples to their respective individual-paired CEC sample, significantly higher levels of CTG18.1 repeat instability were observed in all CEC samples (Wilcoxon test, p=0.004). Furthermore, molecules ranging from 5,439-35,561 bp were only detected in the CEC-derived samples, equating to approximately 1,800-11,900 repeats, with peaks of molecules ranging from 3,900-6,400 repeats (**Figure 2**). In addition, we also investigated three FECD dermal fibroblast lines, generated from CTG18.1 expansion-positive cases, to further explore repeat instability within another unaffected cell type (**Table S1**). When analysed by OGM, CTG18.1 molecule sizes were comparable to leukocytes (mean repeat length ∼97 CTG repeats) (**Figure S1**).

**Figure 2:**
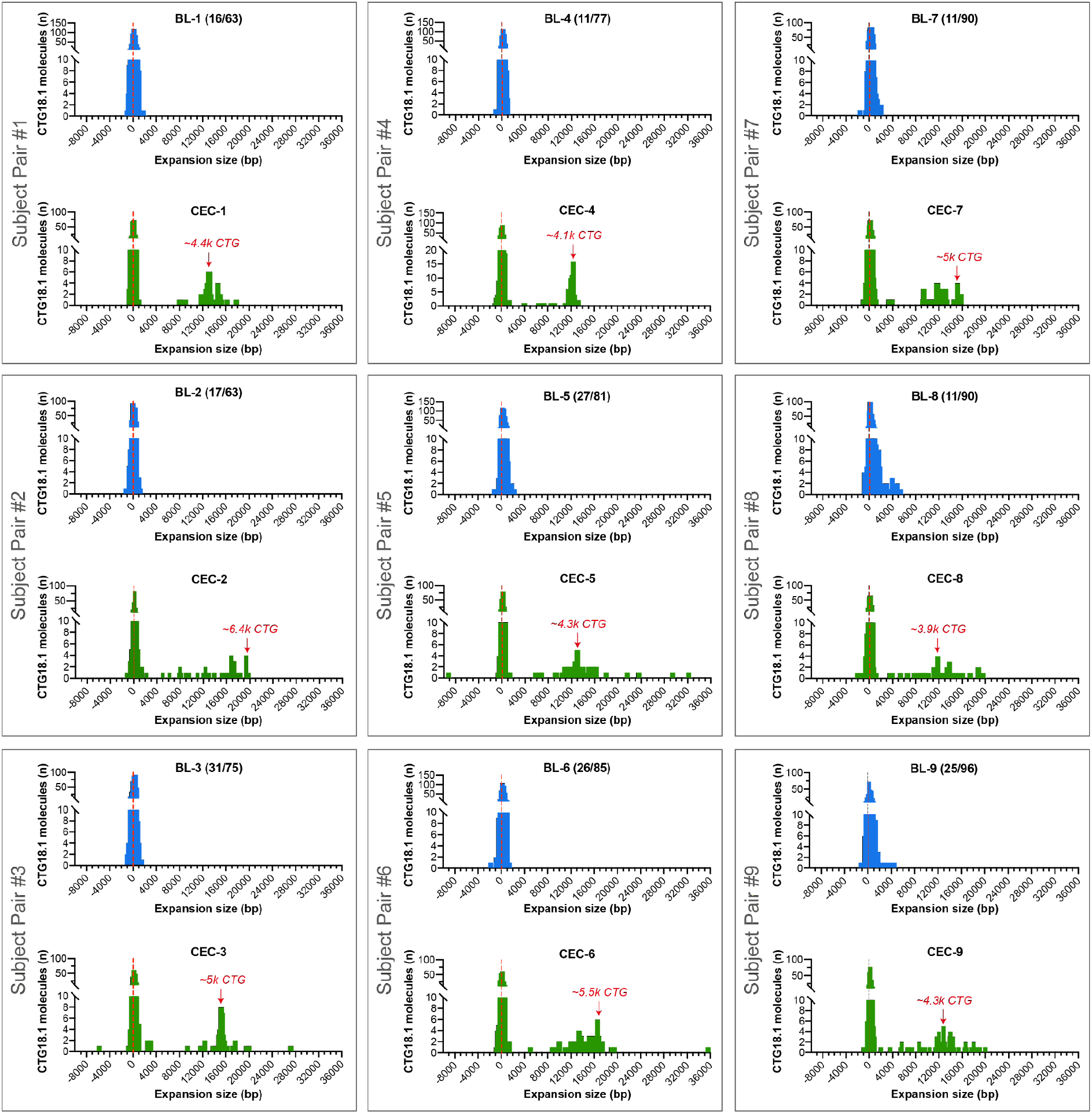
Diseased corneal endothelial cells (CECs) display increased levels of CTG18.1 somatic instability. A series of paired peripheral blood leukocyte-derived (BL1-9 in blue) and corneal endothelial-derived (CEC1-9 in green) gDNA samples from nine unrelated FECD patients analysed by OGM. The size (bp) of the CTG18.1 repeat-containing molecules is plotted (x-axis) against the total number of CTG18.1 molecules detected (y-axis). Red arrows depict the bin with most molecules detected above the 5,439 bp threshold observed exclusively within the CEC-derived gDNA samples.

The vast majority of FECD cases harbour only one expanded allele (*e.g.* all nine cases presented in **Figure 2**). However, we acquired corneal endothelial tissue and subsequently established a primary CEC culture from one additional case with bi-allelic CTG18.1 expansions (baseline CTG18.1 expansion status of 57/93 by STR-analysis of leukocyte-derived DNA). OGM molecule distributions from this sample, alongside representative bi-allelic non-expanded and mono-allelic expanded CEC-derived gDNA FECD samples, are shown in **Figure 3**. Although we could not phase the OGM data, these data illustrate how the zygosity status of the expansion influences molecule distributions. The mean molecule length for the bi-allelic expanded CEC sample (CEC-10) was 16,803bp, equating to approximately 5,600 repeats (**Table 1**). This increased mean molecule length is driven by the shift in the ratio of total molecules ≤2,000bp versus >2,000bp compared to the mono-allelic expanded and expansion-negative FECD CECs. Molecules >2,000bp were exclusively detected in expansion-positive CECs, while these data suggest that the population of molecules ≤2,000bp are predominantly attributed to molecules derived from non-expanded CTG18.1 alleles (**Figure 3; Figure S2**).

**Figure 3:**
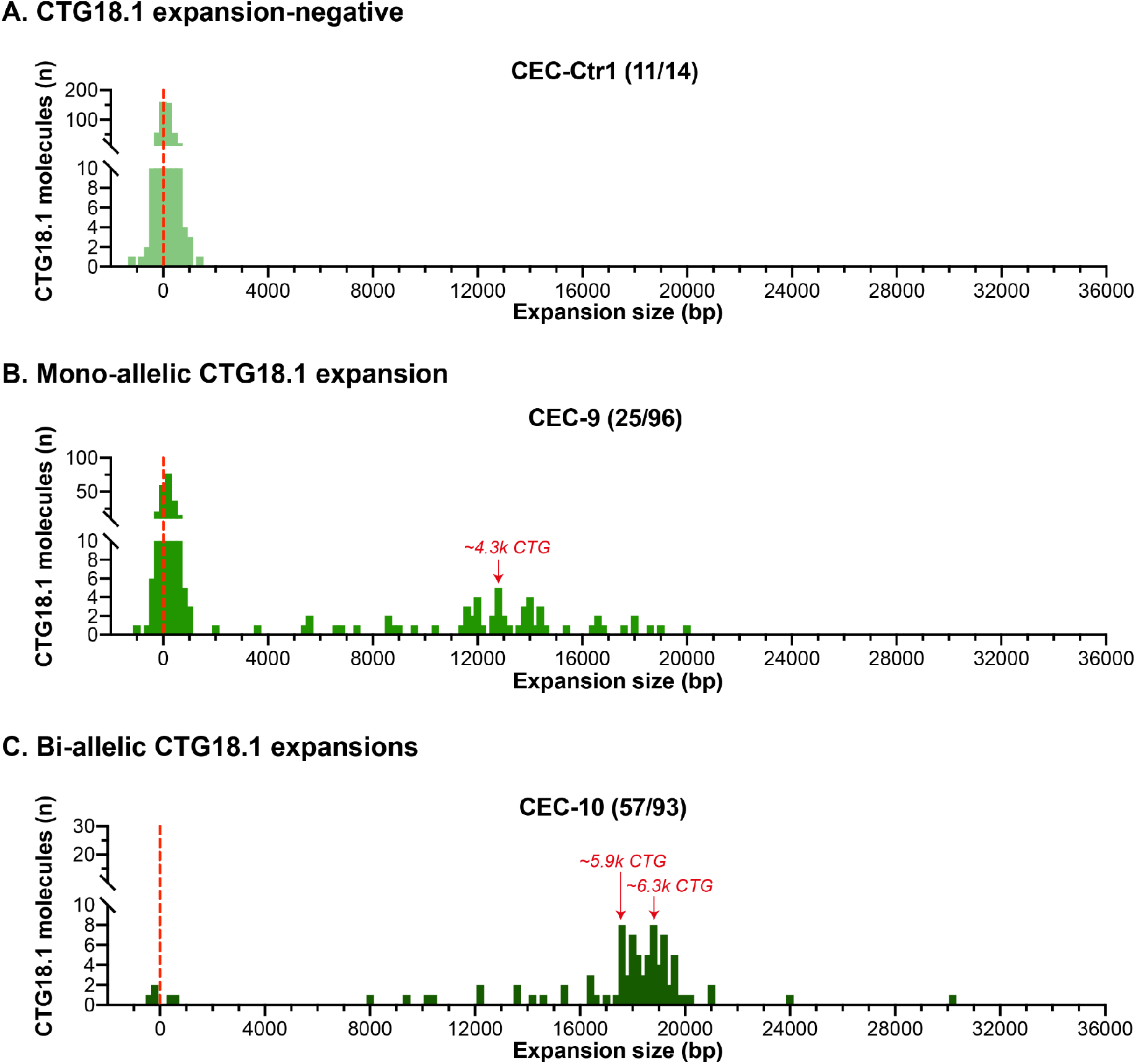
Analysis of bi-allelic CTG18.1 corneal endothelial cells (CECs) indicates that molecules near the detection threshold of optical genome mapping (OGM) represent non-expanded alleles. CEC-derived gDNA samples from three unrelated FECD cases were analysed by OGM. Based on STR-analysis of leukocyte-derived DNA, we classified cases as CTG18.1 **(A)** expansion-negative (CEC-Ctr1), **(B)** mono-allelic expanded (CEC-9) or **(C)** bi-allelic expanded. Baseline STR genotyping results are shown in brackets. Red arrows depict the bin with most molecules detected >2,000 bp threshold, exclusively observed in mono- (CEC-9; B) and bi-allelic (CEC-10; C) expanded CTG18.1 samples.

### Inherited repeat size and cellular context both influence CTG18.1 repeat instability

We have previously demonstrated that expanded CTG18.1 alleles are unstable in leukocytes, an effect that positively correlates with the inherited CTG18.1 allele length (Hafford-Tear et al. 2019). Interestingly, clusters of longer molecules were detected by OGM in samples BL-8 and BL-9, which also had some of the longest inherited repeat sizes detected by STR-PCR within this series (**Figure 2**). Hence, this led us to expand the series of leukocyte gDNA samples to give a more diverse range of inherited expanded repeat sizes (63-107) to further test the capabilities of OGM to detect CTG18.1 somatic instability (**Figure S3**). Molecule counts were plotted by increasing size of CTG18.1 alleles based on the STR genotyping results (**Figure 4A)**. Spearman correlation coefficient analysis revealed a positive correlation between the largest inherited CTG18.1 allele size (determined by STR-PCR of leukocyte gDNA) and the molecule size measured by OGM (0.255, 95% CI: 0.236-0.274) (**Table S2**). In agreement, a simple linear regression on the expansion-positive leukocyte OGM data suggests that the inherited allele size has a strong effect on the detected CTG18.1 molecule size. Given the log transformation of the outcome, the effect of the inherited allele size variable on the OGM measured CTG18.1 molecule size increases exponentially, with increasing inherited allele size (adjusted R^2^ 76.15%, p<0.0001) (**Figure 4B, Table S3**). Adding subject sampling age as an independent variable in the model improved the fit, suggesting that age contributes to the increased levels of somatic instability driven by the inherited allele size (adjusted R^2^ 91.02%, p<0.0001, **Table S3**).

**Figure 4:**
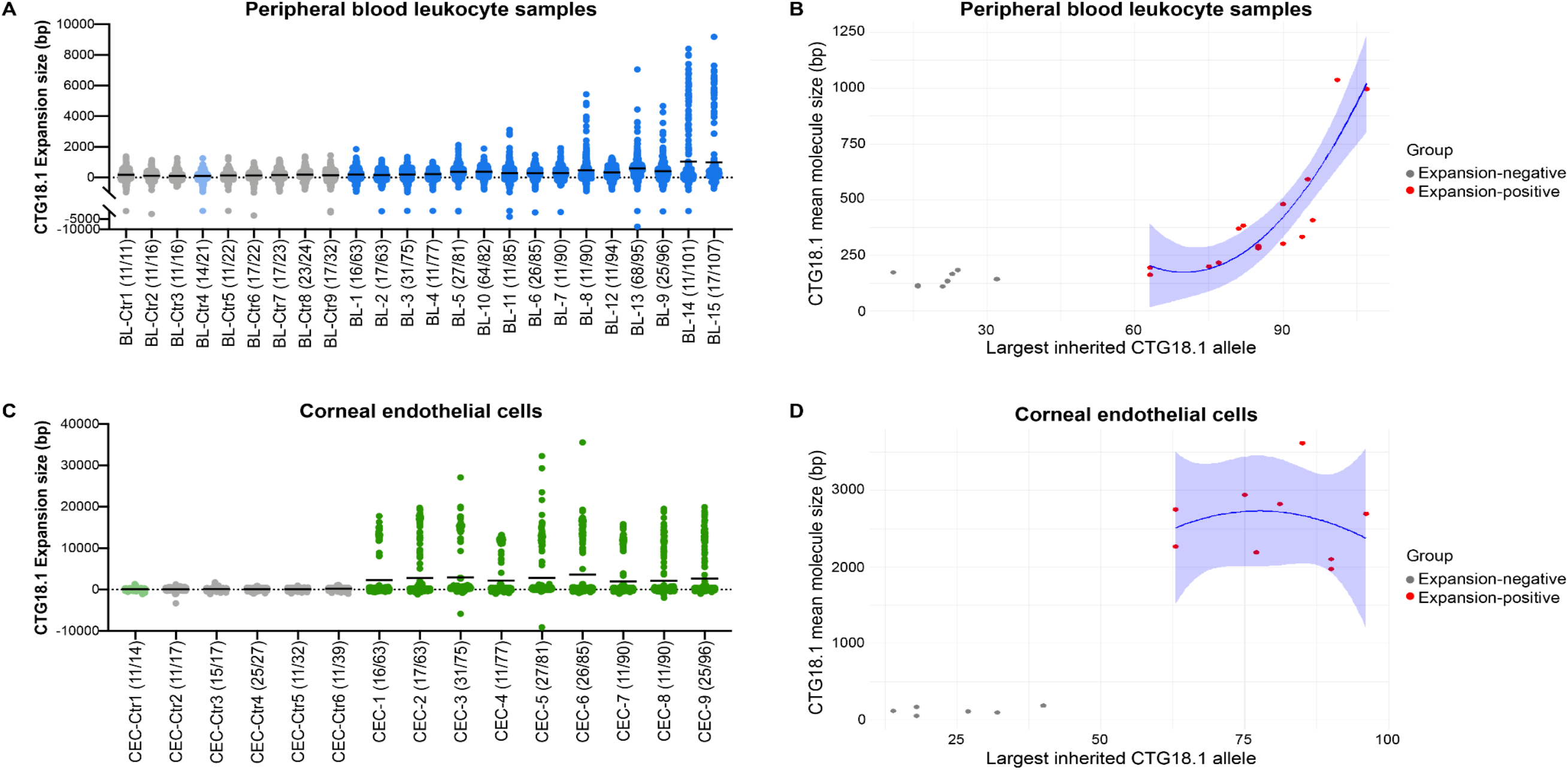
Expanded CTG18.1 alleles behave dynamically in both peripheral blood leukocytes and corneal endothelial cells. Dot plots of molecules detected per individual capturing the distribution of all CTG18.1 alleles analysed in **(A)** peripheral blood leukocytes (BL) and **(C)** corneal endothelial cell (CEC) samples are presented. **(A,C)** Individuals with non-expanded CTG18.1 alleles (<50) are colour-coded in grey (Ctrl). Light blue and light green shades indicate FECD individuals with non-expanded CTG18.1 alleles in their peripheral blood leukocytes or CEC-derived gDNA, respectively. Lines represent the mean CTG18.1 expansion size in base pairs (bp) per sample. **(B,D)** Scatter plots of CTG18.1 mean molecule size (bp) against the largest inherited allele is shown with polynomial regression for the expansion-positive dataset (red) and 95% confidence interval (CI) shown in blue for **(B)** peripheral blood leukocyte and **(D)** CEC sample series. All data series are arranged in order of the largest allele detected per individual according to baseline CTG18.1 genotype determined by the STR-PCR assay and indicated in brackets for each sample.

Next, we wanted to explore if the dynamic and extreme range of CTG18.1 length instability observed in the corneal endothelium is driven by the presence of expanded inherited alleles (≥50 CTG repeats, as detected in peripheral blood leukocyte samples) or if it is a phenomenon observed in all endothelial cells irrespective of their baseline CTG18.1 expansion status **(Figure 4C; Figure S2)**. The corneal endothelium from control tissues was isolated and cultured similarly to the FECD tissues. The control samples included individuals harbouring ≤39 CTG expansions (**Figure 4C**). In addition, we also analysed CECs from an individual with FECD but without CTG18.1 expanded alleles (blood STR genotype 11/14). Within these control samples, we observed an accumulation of molecules around 0 bp, near the detection threshold of the assay, and a complete absence of longer expanded molecules for both control and FECD non-expanded CEC samples (**Figure 4C; Figure S2**). In contrast, remarkable levels of repeat instability were observed for all FECD CEC samples with baseline leukocyte repeat sizes ≥63 determined by STR genotyping. Spearman correlation analysis across the CEC dataset suggested a correlation between the largest inherited CTG18.1 allele size (determined by STR-PCR of leukocyte gDNA) and the molecule size measured by OGM (0.229, 95% CI: 0.201-0.256) (**Table S2).** We found no correlation when we compared this effect within the expansion-negative and the expansion-positive groups in isolation (**Figure 4D; Table S2**). However, when the measured molecule size was compared between the expansion-negative and expansion-positive sample groups, there was a significant difference (Mann Whitney test, p=0.001), suggesting that this is driven by the presence of a CTG18.1 repeat expansion, irrespective of inherited allele size (**Figure 4D**). These data indicate that CTG18.1 repeat instability can be influenced by both the inherited repeat size and cellular context.

## Discussion

FECD is an age-related degenerative eye disease hypothesised to share multiple pathogenic mechanistic parallels with many rare and devastating neurological and neuromuscular diseases also attributed to STR expansions (Fautsch et al. 2021; Malik et al. 2021). Here, for the first time, through the application of OGM, we have shown that CTG18.1 expansions have an exceptionally high level of somatic instability within affected CECs. This finding provides novel mechanistic insights into the pathophysiology of FECD. It also highlights the potential utility of this approach to investigate STR instability more broadly across other repeat loci susceptible to high levels of somatic instability.

A strong correlation between CTG18.1 expansion length in peripheral blood leukocytes and molecular hallmarks of pathology in the corneal endothelium, including the presence of nuclear CUG-specific RNA foci and isoform-specific patterns of *TCF4* downregulation, is well documented (Zarouchlioti et al. 2018; Hu et al. 2018; Powers et al. 2022; Bhattacharyya et al. 2024). Our study has shown that the mean repeat length in mono-allelic expanded CEC lines is 4.4 to 17.0-fold longer than in leukocyte gDNA samples. Specifically, CECs derived from individuals (n=9) with ≥63 baseline CTG18.1 repeats in leukocytes had molecules comprising 1,800-11,900 repeats. Thus, probing CTG18.1 repeat length in these CECs has provided new insights during end-stage corneal endothelial disease and highlights the biological relevance of studying repeat instability directly within primarily affected cells. These data, alongside the low levels repeat of instability and lack of molecular hallmarks in expansion-positive fibroblasts, support the hypothesis that high levels of repeat instability could drive downstream molecular events and further reinforces the utility of primary CECs as an appropriate model to investigate FECD pathophysiology *ex vivo*.

It is now of therapeutic relevance to determine the temporal dynamics of CTG18.1 instability within the corneal endothelium before end-stage disease (i.e., <58 years of age at sampling, reported in this study), given that downstream pathogenic features of CTG18.1 expansion-mediated FECD are hypothesised to be repeat length dependent. Subject sampling age contributed to the increased levels of somatic instability driven by the allele size in the expansion-positive leukocyte-derived samples, in contrast to the expansion-positive CEC subgroup. The lack of correlation in the CECs between subject sampling age and inherited repeat size may be explained by the small sample size in the series or by somatic instability levels already being saturated in the late-stage disease cells.

CTG18.1 molecules with approximately 4,500 repeats were also detected in the immortalised F35T CEC line, previously reported to have 4,500 CTG repeats by Southern blot, suggesting that successive passaging does not grossly alter repeat size. Additionally, *in vitro* studies on CAG repeat instability have reported either a minor (1 CAG repeat increase every 12.4 days) or no correlation between cell proliferation and repeat instability (Gomes-Pereira et al. 2001; Goold et al. 2019). Taken together, these findings suggest that establishing and culturing primary CECs for a single passage, required due to the low cell numbers available from each corneal endothelial explant, is likely to only minimally affect repeat instability and does not explain the magnitude of the repeat instability detected in these cells.

Interestingly, a link between exposure to ultraviolet light and FECD has been suggested in a CTG18.1 expansion-agnostic setting (Liu et al. 2023, 2020). We hypothesise that cumulative UV radiation-inducing DNA damage could contribute to the extreme CTG18.1 instability via the DNA repair-dependent mechanisms in the terminally differentiated CECs. Further studies to explore potential associations between CTG18.1 instability and DNA repair pathways are warranted. Importantly, the extreme levels of instability observed in CTG18.1 expansion-positive CECs were absent from both control and expansion-negative FECD CECs, suggesting that dynamic expansion of the repeat element within the corneal endothelium only occurs when the inherited CTG18.1 repeat size is above a critical threshold (>39 and ≤63 CTG repeats). The presence of RNA foci within CECs has been previously reported in patients with ≥32 CTG18.1 repeats. However, control CECs from a 58-year-old individual harbouring 39 CTG18.1 repeats did not display the high levels of somatic instability observed in the expansion-positive group. This could be explained by differences in the sensitivities of the assays, the contribution of genetic modifiers that may alter the levels of somatic instability, and differences in subject sampling ages. Future efforts will likely further refine the threshold for disease and identify individual modifiers of instability that have been effectively delineated for other STR-associated (Rajagopal et al. 2023).

Analysis of a CEC sample with bi-allelic CTG18.1 expansions suggests that CTG18.1 molecules expand dynamically in the vast majority of CECs with inherited allele size ≥50 CTG repeats at end-stage disease. We anticipate advances in long-read sequencing technologies to enable phasing and sequence-level resolution of the exceptionally large expanded alleles will yield further biological insight into the pathophysiology of CTG18.1-mediated FECD. Notably, the total molecule counts were significantly lower than other samples in the study, suggesting that despite analysing native molecules, OGM is biased against capturing longer molecules.

OGM has an average genome-wide lower detection limit of 500 bp (Zhang et al. 2023). On this basis, we queried its utility for the detection of CTG18.1 expansions in the majority of FECD blood leukocyte-derived gDNA samples, given that 500 bp corresponds to approximately 166 CTG repeats, and in 96.7% of our patient cohort, the expanded repeat sizes ranged from 50-125 CTG repeats (Zarouchlioti et al. 2018). However, analysis of leukocyte samples from individuals with a diverse range of expanded CTG18.1 allele sizes (63-107 inherited repeats) illustrates that OGM can detect an increase in the average CTG18.1 molecule size for inherited alleles ≥63 repeats, but cannot discriminate between samples with alleles ≤32 repeats. We hypothesise this signal is driven by the dynamic instability of CTG18.1, occurring within a fraction of total peripheral blood leukocytes, which increases with an increase in the inherited repeat size. We have previously shown that this dynamic shift in instability occurs at >80 and ≤91 CTG repeats using an amplification-free long-read sequencing approach (Hafford-Tear et al. 2019). The peripheral blood leukocyte gDNA OGM data series presented here is in keeping with this observation, given that the measured molecule size increases exponentially and samples with inherited model repeats ≥81 displayed the most pronounced signature of instability. More generally, these data serve as a benchmark for the wider utility of OGM to explore STR sizing detection and instability thresholds at other loci. To the best of our knowledge, OGM has, to date, only been used to detect and size STR expansions across a limited number of loci and without interrogating the primarily affected tissue (Dai et al. 2020; Efthymiou et al. 2023; Ghorbani et al. 2022; Morato Torres et al. 2022; Facchini et al. 2023). We believe OGM will offer the greatest utility when modal repeat sizes for any given locus exceed the lower detection limit of 500 bp. Nonetheless, our data illustrate that it is also useful when only a small fraction of total DNA molecules exceed the lower detection limit.

In summary, this study illustrates that OGM is a powerful method for detecting and sizing large repeat expansions from native DNA molecules using relatively modest amounts of input material. It also highlights the importance of investigating STR instability in disease-relevant cell types to probe dynamic and tissue-specific mechanisms of somatic instability. Given the exceptionally high levels of CTG18.1 repeat length instability consistently and exclusively observed in affected CECs, these data also lead us to hypothesise that CTG18.1 instability is a key driver of CTG18.1 expansion-mediated disease mechanisms.

## Methods

### Subject recruitment and bio-sample processing

This study was approved by the Research Ethics Committees of University College London (UCL) (22/EE/0090) or the General University Hospital (GUH) Prague (2/19 GACR) and was conducted in accordance/compliance with the Declaration of Helsinki. Written informed consent form was obtained from all participants who provided peripheral blood and/or corneal endothelial samples. The diseased corneal endothelium attached to the Descemet membrane was removed and collected as part of the surgical procedure to alleviate symptoms in individuals with FECD. Control corneal endothelium was similarly isolated from donor corneoscleral discs rejected for clinical transplantation (Miracles in Sight Eye Bank, Texas, USA). Excised tissues were stored in Lebovitz L-15 media (Life Technologies) or Eusol-C (Alchimia) until processing.

### Primary corneal endothelial cell (CEC) culture

CECs were isolated and cultured as described by Zarouchlioti et al. 2018, originally adapted from Peh et al. 2015 (Peh et al. 2015). After excision of the Descemet membrane with the attached endothelium, the tissue was incubated in 0.2% collagenase type I for 2 to 4 hours to digest the Descemet membrane and release the CECs. Following digestion, the CECs were pelleted and resuspended in stabilisation media (Human Endothelial-SFM (Life Technologies) supplemented with 5% foetal bovine serum (FBS), 1% antibiotic/antimycotic, and 10 µM ROCK inhibitor Y-27632 (AdooQ BioSciences). Cells were seeded onto FNC-coated tissue culture plates (Stratech) with stabilisation media. The following day, cell cultures were switched to expansion medium (Ham’s F-12 Nutrient Mix with GlutaMAX Supplement (Life Technologies)/Medium 199 GlutaMAX Supplement (Life Technologies) (1:1), 20 μg/mL ascorbic acid, 1% insulin-transferrin-selenium (Life Technologies), 5% FBS, 1% antibiotic/antimycotic, 10 ng/mL bFGF (Life Technologies) and 10 µM ROCK inhibitor Y-27632 (AdooQ BioSciences). Cells were cultured at 37°C, 5% CO_2_ with media changes twice weekly. Cells were passaged once to expand the cell lines and were collected when they reached maximum confluency.

### F35T cell culture

F35T is an immortalised corneal endothelial line (Saha et al. 2022). F35Ts were cultured on FNC-coated tissue culture vessels with expansion media (recipe described in the previous section). F35T cells were passaged using 1X TrypLE Express (Life Technologies) and the media was changed three times per week. F35Ts were cultured in a controlled environment of 37°C and 5% CO_2_.

### Fibroblast cell culture

Fibroblast cell lines isolated from dermal skin biopsies were cultured in DMEM/F-12, GlutaMAX (Life Technologies) supplemented with 10% FBS and 1% penicillin/streptomycin. Fibroblast cell cultures were maintained at 37°C, 5% CO_2_. Fresh medium was added every 2-3 days and cells were passaged using 1X TrypLE Express (Life Technologies).

### Dual CTG18.1 targeted genotyping

To stratify participants in the study, a dual PCR-based approach was applied to genotype CTG18.1. Genomic DNA was extracted from peripheral blood samples from individuals with FECD (QIAgen Gentra Puregene Blood Kit) or from the scleras of the control corneoscleral discs (QIAgen Blood and Tissue kit) in accordance with the manufacturer’s guidelines.

The CTG18.1 repeat length was determined using a dual PCR-based approach adapted from Weiben E *et al*. 2012 and Warner JP., et al 1996 (Warner et al. 1996; Wieben et al. 2012). In brief, to detect and size CTG18.1 repeat length a STR PCR using a 6’-FAM-conjugated primer (5’-CAGATGAGTTTGGTGTAAGAT-3’) upstream of the repeat and an unlabeled primer (5’-ACAAGCAGAAAGGGGGCTGCAA-3’) downstream. A further triplet-primed (TP) PCR assay to confirm the presence or absence of a CTG18.1 allele above the detection limit of the STR assay (approximately 125 repeats), was also performed utilising a 5’-6-FAM-conjugated primer (5’-AATCCAAACCGCCTTCCAAGT-3’) upstream of the repeat in combination with a reverse primer complementary to the repeat sequence and encompassing a common 5’ sequence (tail) that shares no homology to human genomic sequence (5’-TACGCATCCCAGTTTGAGACGCAGCAGCAGCAGCAG-3’) to serve as an anchor for a second reverse primer (5’-TACGCATCCCAGTTTGAGACG-3’). All PCR amplicons were mixed with the GeneMarker ROX500 ladder (Thermo Fisher Scientific). Capillary electrophoresis was performed on ABI 3730 (Applied Biosystems) and CTG18.1 alleles were sized using GeneMarker (SoftGenetics).

### OGM: sample preparation and cryopreservation and workflow

Fresh blood was collected from all participants before being stored at -80°C. CEC cultures were collected and cryopreserved in 10% DMSO in liquid nitrogen until gDNA extraction. UHMW gDNA was extracted using the “SP or SP-G2 Blood and Cell Culture DNA Isolation” kits (Bionano Genomics) along with the recommended extraction protocols for either frozen blood or cryopreserved cell samples. After homogenisation, the gDNA was labelled with fluorophores at genome-wide DLE-1 recognition motifs, while the DNA backbone was also stained using the “Direct Label and Stain (DLS or DLS-G2)” kits (Bionano Genomics). Stained UHMW DNA molecules were loaded into Saphyre G2.3 or G3.3 chips and single linearised molecules running through the nanochannels were imaged using the Saphyr instrument (Bionano Genomics). High-resolution images were acquired for a throughput of 1.5TB per sample and a minimum expected effective coverage of 400X per sample. As per Bionano’s recommendations, all samples had 14-17 labels/100 kb, >85 map rates, N50≥150 kb ≥230 kb, N50≥20kb ≥150kb.

### Customised OGM analysis pipeline to estimate CTG18.1 repeat length

Molecules were aligned to the hg38 reference using the align_mol_to_ref.py script available in Bionano Solve 3.7.1 software package (https://bionano.com/software-downloads/). CTG18.1 is located between markers 10,414 and 10,415 of chromosome 18 (hg38; chr18:55,584,360-55,594,648). From the alignment files, all molecules overlapping both markers were selected for the downstream analysis pipeline. For each molecule, the expansion size is estimated by calculating the distance difference between the two markers of interest in relation to the reference. The distance between markers 10,414 and 10,415 is 10,288bp. However, a correction was to be applied in this case: when two theoretical binding sites for the labelling enzyme are too close (around 1 kb or less), only one of the two is detected (randomly chosen), meaning that when comparing the molecule with the reference, the average position of these two reference markers must be used. The labelling pattern of our region of interest was also checked in the telomere to telomere hg38, but since it was identical, we proceeded with the correction as shown in **Figure S4**. We applied the same correction scheme used by the standard Bionano DeNovo pipeline: keeping 10,414 (chr18:55,584,360) as the upstream marker and averaging markers 10,415 (chr18:55,594648) and 10,416 (chr18:55,595438) for the downstream marker, which gives a corrected reference distance of 10,683bp. Finally, to correct for the presence of 24 CTG repeats in hg38, we subtracted 72 bp more resulting in a final reference distance of 10,611bp. For the leukocytes and CEC paired series, molecules were plotted based on their size in histograms using 200 bp bin widths. The code to reproduce the analyses described here is available from GitHub (https://github.com/stfacc/extract_gaussian_alleles/blob/main/aln.py).

### Statistical analysis

Spearman correlation analysis was used to investigate the relationship between the largest inherited allele size and the measured molecule size by OGM. Simple and multiple linear regression models using the log-transformed outcome were employed to model the effect of mean inherited allele size per patient on the mean measured molecule size per patient. The model optimisation steps are provided in the results. The final model included an interaction term, and the independent variables were adjusted using mean centring, which eliminated the multicollinearity. We used the Mann-Whitney test to compare the mean measured molecule size per patient between the expansion-positive and expansion-negative subgroups.

## Data Access

The data sets supporting the conclusions of this article are included within the article and its Supplemental Files. All raw CTG18.1 locus-specific molecule OGM data generated in this study will be uploaded and made freely available upon article acceptance.

## Acknowledgements

We thank all affected individuals for participating in this research. We also acknowledge Jana Jedlickova, Monika Pankievic, Natalie Ryskova and Beverly Scott for their technical expertise in DNA extractions and bio-sample processing. We also thank Andrew Powers (Design Therapeutics) for his intellectual input and advice on the application of OGM and Polychronis Kemos for his input on statistical analysis. We would also like to acknowledge Miracles for Sight Eye Bank for providing control corneal tissue for our research.

## Supplemental Data

**Supplemental Figure S1:**
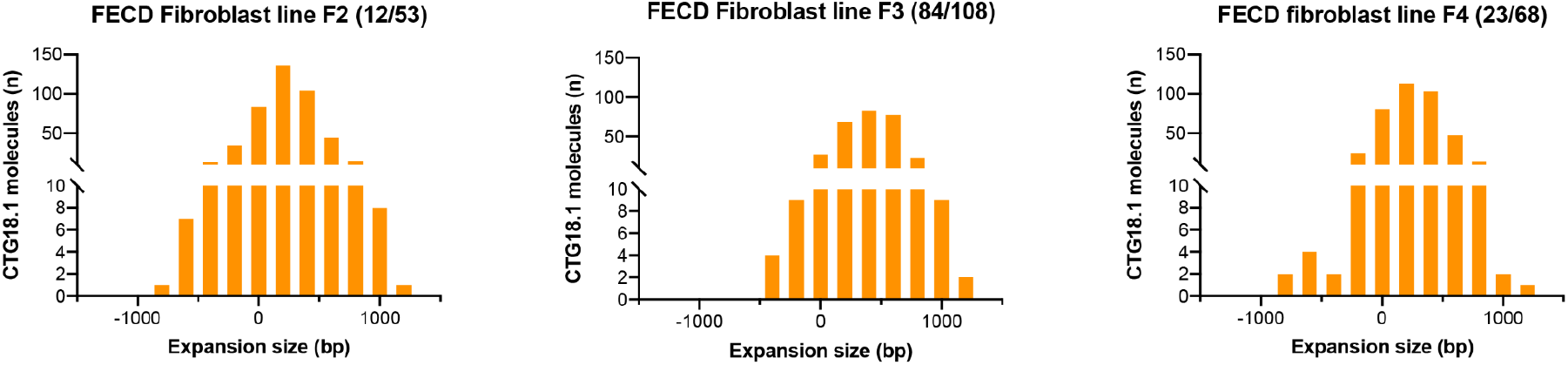
CTG18.1 is somatically stable in expansion-positive fibroblasts. DNA was extracted from cultured dermal fibroblasts isolated from three CTG18.1 expansion-positive individuals with FECD and analysed by optical genome mapping. The size (bp) of the CTG18.1 repeat-containing molecules is plotted (x-axis) against the total number of CTG18.1 molecules detected (y-axis). Baseline CTG18.1 genotypes determined by STR-PCR are shown in brackets for each sample.

**Supplemental Figure S2:**
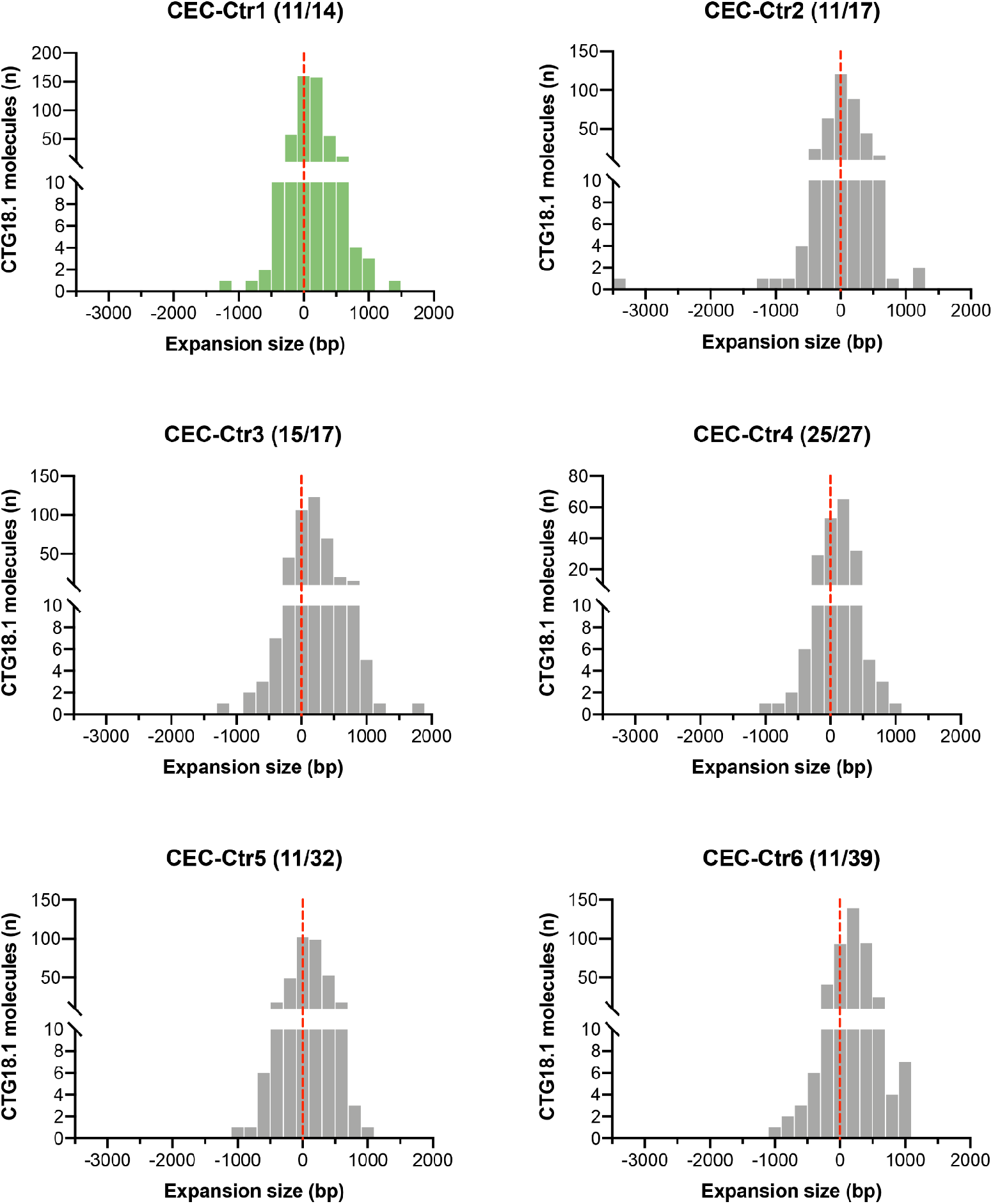
CTG18.1 is somatically stable in control corneal endothelial cells (CECs). DNA was extracted from CECs derived from five healthy control lines and one CTG18.1 expansion-negative FECD-patient derived line and analysed by optical genome mapping (CEC-Ctr1-6). The size (bp) of the CTG18.1 repeat-containing molecules is plotted (x-axis) against the total number of CTG18.1 molecules detected (y-axis). Baseline CTG18.1 genotypes determined by STR-PCR shown in brackets for each sample.

**Supplemental Figure S3:**
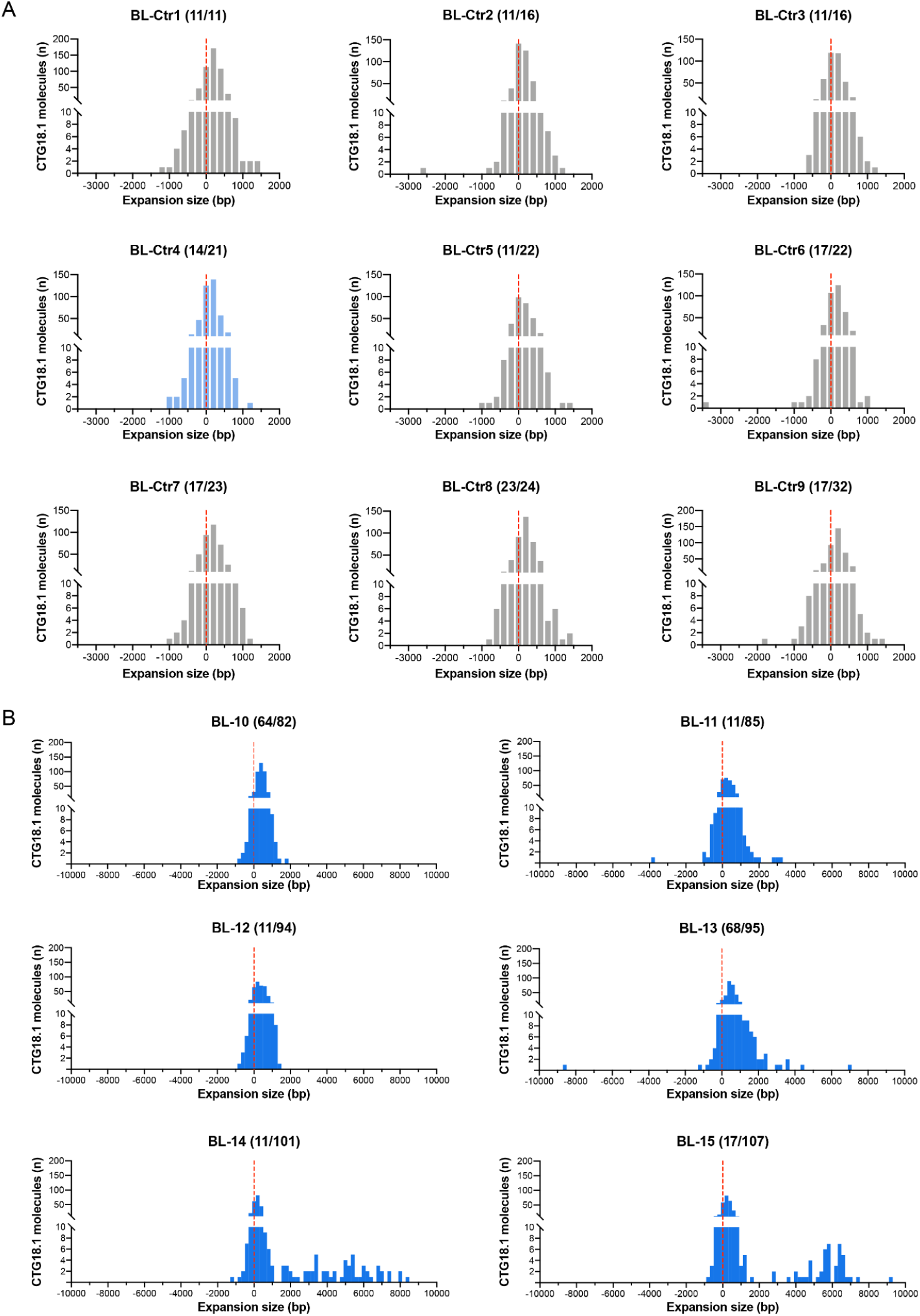
Exploring CTG18.1 instability across a subset of peripheral blood leukocyte samples with a diverse range of repeat sizes. A series of peripheral blood leukocyte-derived gDNA samples from fifteen unrelated individuals were analysed by OGM. (A) Samples BL-Ctr1-9 were derived from eight unaffected (grey) or FECD (light blue) CTG18.1 expansion-negative individuals. (B) Samples BL-10-15 were derived from FECD individuals with mono-allelic CTG18.1 expansions. The size (bp) of the CTG18.1 repeat-containing molecules is plotted (x-axis) against the total number of CTG18.1 molecules detected (y-axis). Baseline CTG18.1 genotypes determined by STR-PCR analysis of leukocyte gDNA are shown in brackets.

**Supplemental Figure S4.**
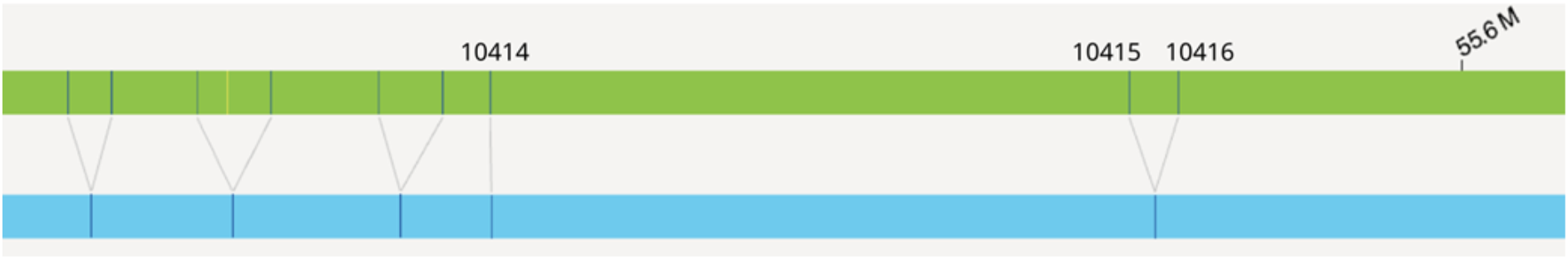
A normal allele (blue) compared with the hg38 reference (green), with matching markers. Several markers, including marker of interest 10415, have more markers in close proximity (<1 kb), however only one of these markers will be detected.

**Table S1:** Optical genome mapping molecule summary of human dermal fibroblasts isolated from CTG18.1 expansion-positive individuals with FECD. a: Fibroblasts isolation and CUG RNA foci data presented in Zarouchlioti et al. 2018. b: *TCF4* isoform RNAScope data presented in Bhattacharyya et al. 2024.

| Subject | Sex | FECD Diagnosis | STR genotype | Subject sampling age | CUG RNA foci | *TCF4* isoform shift | OGM Mean allele length (bp) | OGM Max allele length (bp) | Number of CTG18.1 molecules |
| --- | --- | --- | --- | --- | --- | --- | --- | --- | --- |
| #F2 | Female | ✓ | 12/53 | 71 | No<sup>a</sup> | No<sup>b</sup> | 224 | 1,127 | 446 |
| #F3 | Female | ✓ | 84/108 | 65 | No<sup>a</sup> | No<sup>b</sup> | 396 | 1,225 | 303 |
| #F4 | Female | ✓ | 23/68 | 78 | No<sup>a</sup> | No<sup>b</sup> | 249 | 1,129 | 392 |

**Table S2:**
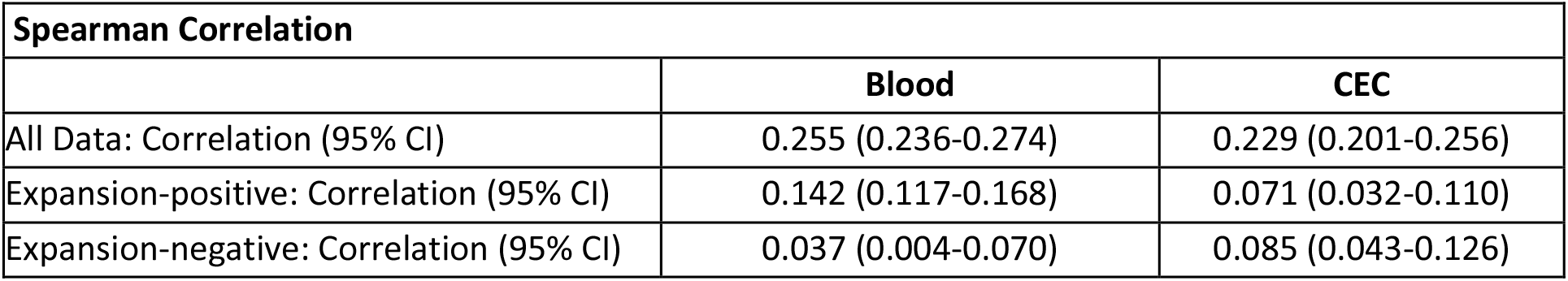
Spearman correlation coefficient analysis exploring the relationship between the largest inherited CTG18.1 allele and the total molecule sizes measured by OGM. Cases with ≥50 CTG repeats (STR analysis of leukocyte-derived DNA) on at least one allele were classified as expansion-positive.

**Table S3:** Simple and multiple linear regression models using the log-transformed outcome to model the effect of mean inherited allele size (determined by STR-analysis of leukocyte-derived DNA) and/or subject sampling age per patient on the mean molecule size measured by optical genome mapping per patient. NA: not applicable.

| **Linear Regression Modelling Results** |  |  |  |  |
| --- | --- | --- | --- | --- |
| Model specification | Adjusted R<sup>2</sup> | p-values for coefficients |  |  |
|  |  | Largest inherited CTG18.1 allele size | Subject sampling age | Interaction |
| Largest inherited CTG18.1 allele size only | 76.15% | <0.0001 | NA | NA |
| Subject sampling age only | 0% | NA | 0.59 | NA |
| Largest inherited CTG18.1 allele and subject sampling age (no interaction) | 79.84% | <0.0001 | 0.10 | NA |
| Largest inherited CTG18.1 allele and subject sampling age (with interaction) | 91.02% | <0.0001 | 0.02 | 0.02 |

